# Esrra regulates Rplp1-mediated translation of lysosome proteins suppressed in non-alcoholic steatohepatitis and reversed by alternate day fasting

**DOI:** 10.1101/2021.11.16.468891

**Authors:** Madhulika Tripathi, Karine Gauthier, Reddemma Sandireddy, Jin Zhou, Priyanka Gupta, Suganya Sakthivel, Wei Wen Teo, Yadanar Than Naing, Kabilesh Arul, Keziah Tikno, Sung-Hee Park, Yajun Wu, Lijin Wang, Boon-Huat Bay, Lei Sun, Vincent Giguere, Pierce K. H. Chow, Sujoy Ghosh, Donald P. McDonnell, Paul M. Yen, Brijesh K. Singh

## Abstract

**Background:** Currently, little is known about the mechanism(s) regulating global and specific protein translation during non-alcoholic steatohepatitis (NASH).

**Methods:** We used puromycin-labelling, polysome profiling, ChIPseq and ChIP-qPCR, and gene manipulation *in vitro* and in dietary mouse models of NASH in this study.

**Results:** Using unbiased label-free quantitative proteome, puromycin-labelling and polysome profiling, we observed a global decrease in protein translation during lipotoxicity in human primary hepatocytes, mouse hepatic AML12 cells, and livers from a dietary mouse model of NASH. Interestingly, proteomic analysis showed that Rplp1, which regulates ribosome and translation pathways, was one of the most downregulated proteins. Moreover, decreased Esrra expression and binding to the Rplp1 promoter, diminished Rplp1 gene expression during lipotoxicity. This, in turn, reduced global protein translation and Esrra/Rplp1-dependent translation of lysosome (Lamp2, Ctsd) and autophagy (sqstm1, Map1lc3b) proteins. Of note, Esrra did not increase its binding to these gene promoters or their gene transcription, confirming its regulation of their translation during lipotoxicity. Notably, hepatic Esrra-Rplp1-dependent translation of lysosomal and autophagy proteins also was impaired in NASH patients and liver-specific *Esrra* knockout mice.

Remarkably, alternate day fasting induced Essra-Rplp1-dependent expression of lysosomal proteins, restored autophagy, and reduced lipotoxicity, inflammation, and fibrosis in hepatic cell culture and *in vivo* models of NASH.

**Conclusion:** Esrra regulation of Rplp1-mediated translation of lysosome / autolysosome proteins was downregulated during NASH. Alternate day fasting activated this novel pathway and improved NASH, suggesting that Esrra and Rplp1 may serve as therapeutic targets for NASH. Our findings also provided the first example of a nuclear hormone receptor, Esrra, to not only regulate transcription but also protein translation, via induction of Rplp1.

## 1. Introduction

Non-alcoholic fatty liver disease (NAFLD) is a common and chronic liver disease that has emerged as a significant public health concern worldwide [1]. NAFLD comprises a range of liver disorders, from the excessive buildup of fat in hepatocytes, known as steatosis, to a more severe condition known as nonalcoholic steatohepatitis (NASH) [2, 3]. NASH has the potential to advance into cirrhosis and hepatocellular carcinoma, and is the leading cause for liver transplantation [4, 5]. NASH also is closely linked to metabolic disorders such as obesity, insulin resistance, and dyslipidemia [6, 7]. Currently, limitations in our understanding of the metabolic and molecular defects in NASH have prevented the development of effective strategies to diagnose and treat NASH.

We and others have shown that autophagy [8, 9] is dysregulated in NASH which can lead to defects in fatty acid metabolism, inflammation, and fibrosis [10–12]. Restoration of hepatic autophagy reduced NASH and delayed its progression [11, 13–16]. The mechanism for reduced autophagy in NASH is not known but is associated with lipotoxicity, a condition where excessive intracellular lipids generate reactive oxygen species and ER stress [8] [6]. During ER stress, the unfolded protein response (UPR) drives expression of transcription factors such as ATF4/6, XBP, IRE1α, and phosphorylation of eIF2α to attenuate global translation. Interestingly, higher eIF6 level marks the progression from NAFLD to HCC, independent from other translation machinery factors [17]. Currently, little is known about the relationships among autophagy, lipotoxicity, and protein translation during NASH.

The expression of nuclear receptors such as thyroid hormone and peroxisome proliferator-activated receptors (TRs, PPARs) are reduced in NASH [9, 18–21]. These receptors also regulate the expression of autophagy genes. During NASH, the expression of DIO1, the enzyme that converts T4 to T3 also is reduced and there is decreased intrahepatic T3 concentration in NASH to decrease autophagy and lipid metabolism [22]. Another nuclear receptor, estrogen-related receptor alpha (Esrra/ERRα) is an orphan nuclear receptor whose transcriptional activity is dependent upon heterodimerization with Ppar *gamma* coactivator-1 *alpha* (Ppargc1a/Pgc1α) [23]. Although Esrra has been recognized as a transcription factor for mitochondrial genes regulating mitochondrial activity, biogenesis, and turnover by mitophagy [23], its role(s) in the regulation of ribosomes and the translation of lysosome/autophagy proteins has not been described previously. In this report, we identified a novel Esrra-Rplp1 pathway that regulated protein translation of autophagy/lysosome proteins. This pathway was impaired in NASH and could be stimulated by Esrra overexpression or alternate day fasting. Our studies also showed unexpectedly for the first time that a nuclear receptor not only regulated transcription but was able to modulate the translation of a specific subset of autophagy/lysosome proteins.

## 2. Results Land Discussion

### 2.1 Supressed ribosome, protein translation, translation-associated pathways and mitophagy were observed in NASH

Previous studies have linked ER-stress and specific translation factors (eIF6, eIF2, RPS6KB1) with the increased translation of lipid biosynthesis, cytokines, and inflammatory proteins during NASH pathogenesis [17, 24–26]. However, the regulation of global translation during NASH is not well understood. To understand this, we employed liver tissues collected from mice fed a Western diet with fructose (WDF) (**Figure 1A**) that mimicked NASH progression in humans and previously characterized by us [2, 3, 7]. Hematoxylin & Eosin (H&E) staining of these liver tissues showed NASH features (*e.g.,* micro/macro steatosis, hepatocyte hypertrophy/ballooning and inflammatory foci) in mice fed WDF for 16 weeks whereas only steatosis was found in mice fed WDF for 8 weeks (**Figure 1B).** Liver triglyceride (TG) was also progressively increased from 8 and 16 weeks (**Figure 1C)**. Hepatic inflammatory and fibrosis gene expression was significantly increased in mice fed WDF for 8 weeks and further increased in those fed WDF for 16 weeks (**Figure 1D**). To understand protein translation in NASH, we performed polysome profiling in the liver tissues from mice fed WDF or NCD for 16 weeks and found that polysome fractions were reduced while the 80S fraction was increased in livers from the former (**Supplementary Figure 1A**), demonstrating that global hepatic translation was suppressed during NASH. To better understand the global changes in proteome, we performed an unbiased Label-Free Quantitative (LFQ) proteomics analysis in liver tissues from mice fed WDF 16 weeks or NCD. First, we analyzed the pathways associated with the upregulated (≥1.5FC) proteins using DiGeNET disease discovery platform, Kyoto Encyclopedia of Genes and Genomes (KEGG) 2021 and Reactome 2022 pathway databases (**Figures 1E and F**; **Supplementary Table 1**). Among the diseases discovered from DisGeNET (adjusted p value < 0.05), we observed the upregulated proteome corresponded with non-alcoholic fatty liver disease, steatohepatitis, liver cirrhosis, abnormal liver enzymes and transaminases, obesity, impaired glucose tolerance, insulin resistance, hyperlipidemia, hypertriglyceridemia, and metabolic disease phenotypes. Among the pathways analyzed from KEGG and

**Fig. 1.**
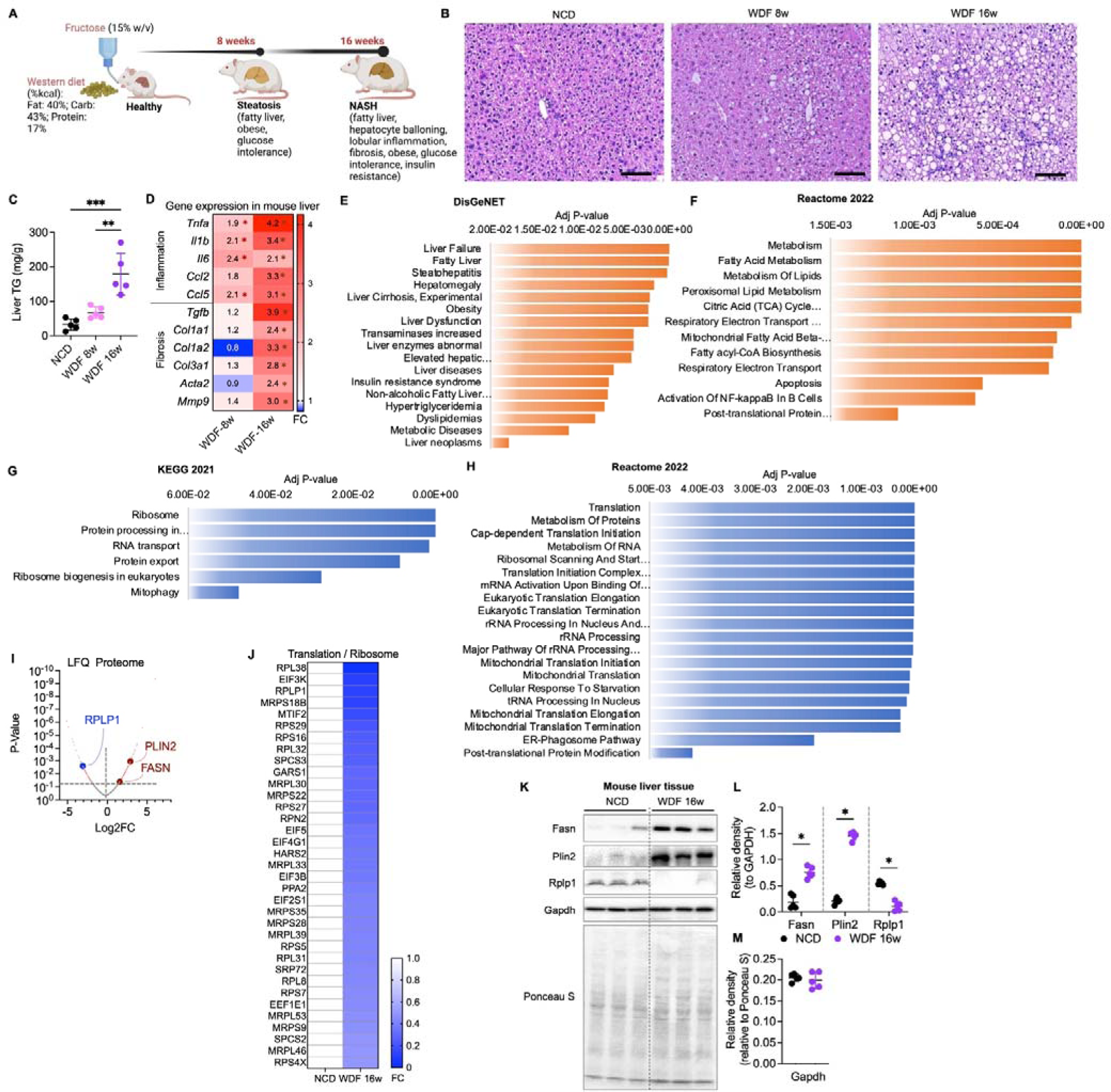
Ribosome, translation and translation-associated pathways were suppressed in WDF fed mice livers. (A) Illustration represents NASH progression in mice fed WDF for 8 and 16 weeks, and characterized by Tripathi et. al. [2]. (B) Representative micrograph of H&E staining in liver tissues collected from mice fed Western diet and fructose (WDF) for 8 and 16 weeks to generate steatosis and NASH and compared to controls fed normal chow diet (NCD) (n=5 per group). Scale bars are 100 μM. (C) Triglycerides (TG) measurement in the liver tissues. (D) Heat map representing RT-qPCR analysis of inflammatory and fibrosis genes in the liver tissues from mice fed WDF diet for 8 or 16 weeks (n=5 per group). Gene expression is normalized to Gapdh. (E and F) DisGeNET and Reactome 2022 pathway analysis of upregulated (≥1.5FC) proteins measured using label-free quantitative (LFQ) proteomics in liver tissue of mice fed WDF for 16 weeks when compared to NCD. (G and H) KEGG 2021 and Reactome 2022 pathway analysis of downregulated (≤0.5FC) proteins. (I) Valcano plots represents significantly downregulated (RPLP1 among others) and upregulated (PLIN2 and FASN among others) proteins. (J) Heat map showing proteins’ expression (Fold Change, FC) regulating Ribosome and Translation pathways shown in panels G and H. (K) Representative Western blots of livers from mice fed NCD or WDF diet fed 16 weeks (n=5 per group). Ponceau S-stained membrane is representing the protein loading. (L) Plots represent relative density of corresponding Western blots normalized to Gapdh. (M) Plots represent relative density of corresponding Western blots normalized to Ponceau S. Ponceau S-stained membrane is consistent with the Gapdh expression. Levels of significance: *P<0.05; **P<0.01; ***P<0.001; ****P<0.0001.

Reactome (adjusted p value < 0.05), we found a significant association with the activation of fatty acid metabolism, lipid acyl-CoA biosynthesis, TCA, fatty acid oxidation, apoptosis, and NF-*kappa*B, biosynthesis of unsaturated fatty acids, cholesterol metabolism, glucagon signaling, insulin signaling, insulin resistance, Hedgehog ligand biogenesis pathways, ChREBP, RUNX2, NOTCH4, MAPK, PPAR pathways, Interleukin-1 family signaling, and cellular senescence pathways. These discovered diseases and upregulated cellular pathways further validated NASH model in an unbiased manner [6, 12, 27–31]. To our surprise, downregulated proteins (≤ 0.5 FC) were highly associated with translation (adjusted p value < 1.88E-15), metabolism of amino acids and proteins (adjusted p value < 5.56E-13), ribosome (adjusted p value < 1.88E-15), metabolism of RNA (adjusted p value < 5.79E-09), and protein processing, RNA transport, ribosomal biogenesis, mitophagy, ER-phagosome, with other translation-associated pathways to a significant levels (adjusted p value < 0.05) (**Figures 1G and H**; **Supplementary Table 1**). Further analysis of ribosome-associated proteins identified RPL38, RPLP1, RPS29 and RPS16 among the top ten downregulated proteins (**Figures 1I and J**). PLIN2 and FASN were also identified as significantly upregulated proteins involved in both fatty acid metabolism and fatty acyl-CoA biosynthesis (**Figures 1F**). We confirmed that the expression of both Plin2 and Fasn were increased while Rplp1 was decreased significantly in the liver tissues from mice fed WDF for 16 weeks suggesting the latter was not involved in their translation (**Figures 1K and L**).

Our findings indicated that although overall translation was suppressed, the translation of certain proteins was distinctly regulated. The translation of Gapdh remained consistent when compared with Ponceau S staining of blotted proteins (**Figure 1K and M**), making it a reliable loading control for subsequent data evaluation. Past research also has documented GAPDH protein aggregation, its migration to the nucleus, and diminished enzymatic function in liver diseases with no alterations in overall protein expression [32, 33].

### 2.2 Lipotoxic condition downregulated Esrra-Rplp1 axis and impaired protein translation of lysosome/autophagy proteins

We and others previously have demonstrated that Esrra transcriptionally regulates autophagy and mitophagy [23, 34–36]. However, its role in the regulation of translation remains unclear. Hepatic Esrra ChIP-seq data revealed that Esrra binds to its own promoter (**Figure 2A**), suggesting its transcription was involved in a positive-feedback loop [23, 37]. Notably, Esrra was found to bind specifically to the *Rplp1* gene’s promoter (**Figure 2A**), which also happened to be among the top ten most downregulated proteins involved in ribosome and translation pathways during NASH (**Figures 1H and I**). Of note, Esrra was not observed to bind to any of the other ribosomal gene promoters ranked among the top ten (**Figures 1J**). We next performed a ChIP-qPCR analysis on AML12 cells treated with and without the saturated fatty acid, palmitic acid (PA; 0.5 mM for 24 h) [6, 14, 29, 38, 39]. We observed decreased recruitment of Esrra on the ERRE (estrogen-related receptor response element) and Polr2a on the TATA box of the *Esrra* gene promoter during lipotoxic condition (**Figure 2B**). Similarly, recruitment of Esrra on the ERRE and Polr2a on the TATA box of the *Rplp1* gene promoter was reduced by PA treatment in AML12 cells (**Figure 2C**). Moreover, these effects were consistent with reduced *Esrra* and *Rplp1* gene expression observed in AML12 cells treated with PA (**Figures 2D and E**).

**Fig. 2.**
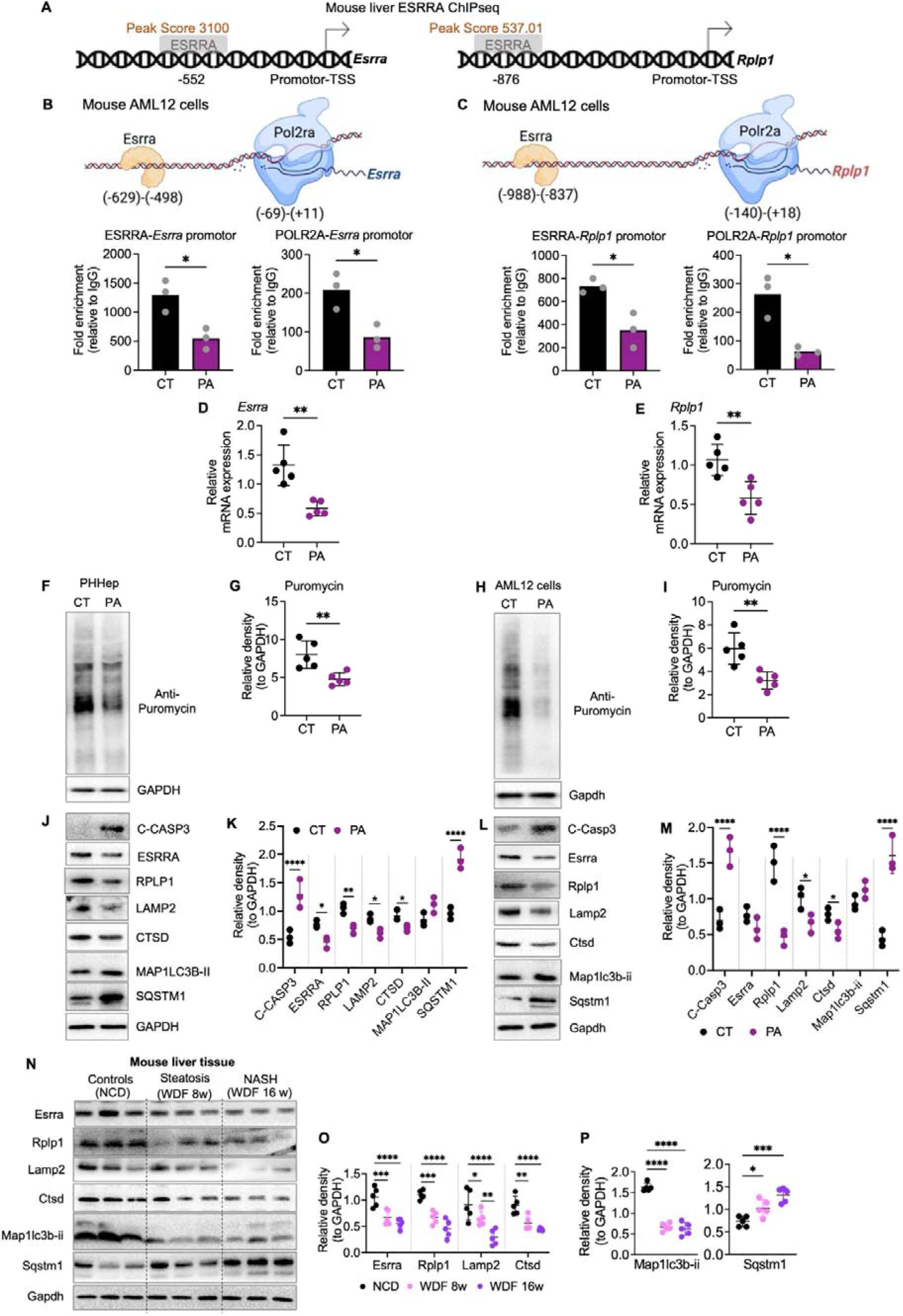
Protein translation was decreased in lipotoxicity, and regulated by Esrra-Rplp1 axis. (A) Cartoon is representative of hepatic Esrra binding on *Esrra* or *Rplp1* promotor region with peaks score and at the site as indicated. The mouse liver ChIPseq data was published elsewhere [37]. (B and C) ChIP-qPCR analysis of Esrra and Polr2a binding on both *Esrra* and *Rplp1* gene promoter near ERRE and TATA box regions respectively at indicated sites. (D and E) RT-qPCR analysis of relative gene expression in PA treated AML12 (n=5 per group). Gene expression is normalized to Gapdh. (F) Representative Western blots of 24 h PA (0.5 mM) treated primary human hepatocytes (PHHep; n=3-5 per group) to analyze puromycin-labeled proteins as a measure to translation. (G and H) Plots represent relative density of corresponding Western blots normalized to Gapdh. (I) Representative Western blots of 24 h PA (0.5 mM) treated AML12 cells (n=3-5 per group). (J and K) Plots represent relative density of corresponding Western blots normalized to Gapdh. (L) Representative Western blots of livers from mice fed NCD or WDF diet (n=5 per group). (M and N) Plots represent relative density of corresponding Western blots normalized to Gapdh. Levels of significance: *P<0.05; **P<0.01; ***P<0.001; ****P<0.0001.

To further understand the role of Esrra on Rplp1 expression and its potential regulation of autophagy and lysosome proteins, we treated primary human hepatocytes (PHHep) and mouse hepatic AML12 cell line with PA (0.5 mM for 24 h). Further, to ascertain protein translation rates, we briefly incubated the PA-treated or untreated cells with puromycin (20 μg/ml) for 15 minutes just prior to protein extraction using a standard assay for measuring protein translation activity [40–44]. Puromycin is a natural aminonucleoside resembling the 3′ end of aminoacylated tRNAs that integrates into the C-terminus of growing nascent protein chains during ribosome-mediated protein translation [40]. These puromycin-labeled proteins then were analyzed by Western blot analysis.

We observed significant decreases in the puromycin-labeled proteins in both PHHep and AML12 cells after PA treatment (**Figure 2F-I)**, suggesting there was decreased protein translation activity. We also confirmed the presence of lipotoxicity in PA-treated PHHep and mouse hepatic AML12 cells by demonstrating increased expression of inflammation (*Tnfa, Il1b, I6, Ccl2,* and *Ccl5*) and fibrosis (*Tgfb, Col1a1, Col1a2, Col3a1, Acta2* and *Mmp9*) genes, and Casp3 cleavage (a hallmark of lipotoxicity [6]) (**Figure 2J-M; Supplementary Figure 2A**).

We also found the mRNA and protein expressions of Esrra and Rplp1 were downregulated in PHHep and AML12 cells treated with PA (**Figures 2J-M**), confirming these proteins were mainly transcriptionally regulated (**Figures 2D and E**). Next, to examine whether lysosome and autophagy proteins might be affected by Esrra and Rplp1 downregulation during lipotoxic conditions, we analyzed the protein expression of Lamp2, Ctsd, Map1lc3b, and Sqstm1. Indeed, Lamp2, Ctsd, and Map1lc3b-ii protein expression decreased, whereas their mRNA expression did not significantly change with PA treatment (**Figures 2J-M; Supplementary Figure 2B**). Further, Sqstm1 protein expression increased while its mRNA significantly decreased (**Figures 2J-M; Supplementary Figure 2B**). These findings were consistent with a late block in autophagy caused the accumulation of Sqstm1 protein as reported previously [23]. Thus, lipotoxic conditions were sufficient to down-regulate the Esrra-Rplp1-lysosome pathway and decrease autophagy flux.

We confirmed these *in vitro* findings in liver tissues from mice fed WDF for 8 and 16 weeks (**Figures 1A).** Hepatic Esrra, Rplp1, Lamp2, and Ctsd protein expression decreased during progression from steatosis (WDF 8w) to NASH (WDF 16w) in mice fed WDF (**Figures 2N and O)**. Hepatic Map1lc3b-ii also significantly decreased whereas Sqstm1 significantly accumulated, suggesting autophagy block in the livers of mice fed WDF (**Figures 2N and P**).

To determine whether the decreased expression of Esrra, Rplp1, lysosome and autophagy proteins was due to reduced translation activity of these proteins, we performed immunoprecipitation of puromycin-labeled proteins and Western blotting to identify newly translated Esrra, Rplp1, Lamp2, Ctsd, Map1lc3b-ii, and Sqstm1 proteins in AML12 cells treated with PA. There was decreased puromycin-labeling / translation activity of Esrra, Rplp1, Lamp2, Ctsd, Map1lc3b, and Sqstm1 proteins (**Figures 3A and B**), although total protein expression of the latter was increased in the input lysate due to autophagy block (**Figures 2J-M, 3A and B**). To demonstrate the selective decrease in translation of these proteins, we also found that puromycin-labeling of NASH-associated protein Plin2 increased during fat droplet formation and lipotoxicity (**Figures 3C**). Taken together, our findings showed that decreased transcription and translation of Esrra and Rplp1 during lipotoxicity selectively reduced translation of lysosome-autophagy proteins to decrease autophagy, a key defect in the NASH pathogenesis [2, 9, 11, 15, 45].

**Fig. 3.**
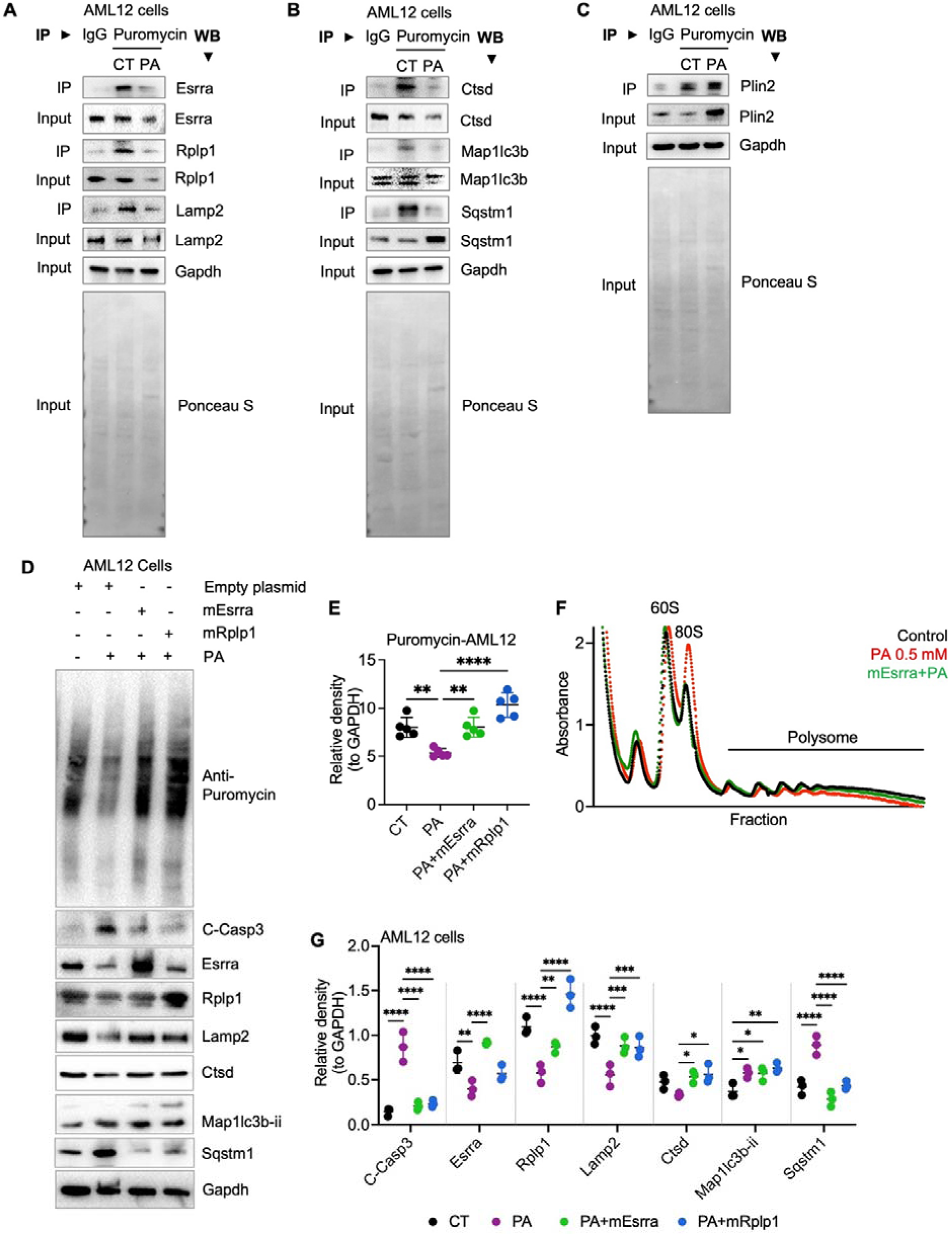
Esrra or Rplp1 overexpression increased translation and lysosome-autophagy protein expression and activity in hepatic cells. (A, B and C) Western blots of immunoprecipitated (IP) proteins labelled by puromycin or IgG in 24 h PA (0.5 mM) treated AML12 cells to analyze puromycin-labeled proteins as a measure to recent translation events. (D) Representative Western blots of 24 h PA (0.5 mM) treated AML12 cells overexpressed with Esrra or Rplp1 or empty vector. (E and F) Plots represent relative density of corresponding Western blots normalized to Gapdh. (G) Representative polysome profile in BSA (control) or PA treated (0.5 mM, 24 h) AML12 cells with or without Esrra overexpression. Levels of significance: *P<0.05; **P<0.01; ***P<0.001; ****P<0.0001.

### 2.3 Activating Esrra-Rplp1-dependent translation of lysosome/autophagy proteins decreased lipotoxicity in vitro

Induction of lysosome-autophagy activity was beneficial for NASH and several other metabolic diseases [9, 15, 16, 46, 47]. Accordingly, we next examined whether activating Esrra-Rplp1-lysosome protein translation pathway could activate autophagy by overexpressing Esrra and Rplp1 in AML12 cells and treated with PA. Either Esrra or Rplp1 overexpression increased the overall puromycin-labeled protein expression and restored ribosome-dependent protein translation that was decreased in PA-treated cells (**Figures 3D and E**). Using polysome profiling, we found that PA substantially increased 80S ribosome fraction and decreased polysome fractions to demonstrate inhibition of mRNA translation (**Figure 3F**). However, Esrra overexpression during PA treatment decreased 80S ribosome fraction and increased polysome fractions similar to control levels to show there now was restoration of efficient mRNA translation elongation. The data suggest that activating Esrra stimulated ribosome-dependent protein translation in PA-treated cells. Furthermore, Esrra or Rplp1 overexpression also decreased Casp3 cleavage in PA-treated AML12 and restored the expression of Lamp2 and Ctsd proteins (**Figure 3D and G**). Moreover, increased Map1lc3b-ii and decreased Sqstm1 proteins in Esrra or Rplp1 overexpressed AML12 cells treated with PA, suggesting there was improved autophagy (**Figures 3D and G**). Additionally, the induction of inflammatory and fibrosis genes expression by PA was attenuated by Esrra and Rplp1 overexpression (**Supplementary Figure 3**).

### 2.4 Esrra-Rplp1-lysosome protein translation pathway was impaired in the liver tissues of NASH patients and mice

Next, we analyzed liver tissues collected from patients with hepatosteatosis (n=7), NASH (n=9), and control (n=11) based upon liver histology (**Supplementary Table 2**). There was increased expression of inflammatory (*TNFA, IL1B, IL6, CCL2 and CCL5*) and fibrosis (*TGFB, COL1A1, COL1A2, COL3A1, ACTA2* and *MMP9*) genes in livers from patients with hepatosteatosis when compared to controls that was further increased in livers from patients with NASH (**Supplementary Figure 4A**). These findings were consistent with the histological classification of the liver tissues. Remarkably, we found that *ESRRA* and *RPLP1* gene and protein expression were significantly decreased in these NASH liver tissues. Additionally, LAMP2, CTSD, and MAPLC3B-II protein expression decreased in hepatosteatosis and further declined in NASH (**Supplementary Figures 4B and C**) whereas SQSTM1 protein significantly accumulated in NASH consistent with late autophagy blockage.

To determine whether decreased hepatic Esrra, Rplp1, Lamp2, and Ctsd proteins’ expression and autophagy block in NASH were associated with impaired translation *in vivo*, we fed mice with methionine-choline deficient (MCD) diet for 3 and 6 weeks to rapidly generate NASH [38, 39], and labeled the proteins with puromycin (*i. p.*, 20 mg/Kg body weight) just 30 min before euthanasia. Histological analysis using H&E staining showed NASH progression between 3 and 6 weeks in mice fed MCD diet *i.e.* steatosis, inflammatory foci and fibrotic regions (**Figure 4A).** These mice also exhibited progressive increases in liver triglycerides (TG), and inflammatory/fibrosis gene expression at 3 and 6 weeks (**Figures 4B and C**). Moreover, we observed progressive decreases in puromycin-labeled proteins in conjunction with decreased expression of Esrra, Rplp1, Lamp2, Ctsd and Map1lc3b-ii proteins and increased expression of Sqstm1 protein in mice fed MCD diet for 3 and 6 weeks, respectively (**Figures 4D-F**). Similar to the results found in PA-treated AML12 cells (**Figure 3A-C**), there was decreased puromycin-labeling (*i.e.* decreased translation activity) of Esrra, Rplp1, Lamp2, Ctsd, Map1lc3b-ii, and Sqstm1 proteins, even though Sqstm1 protein expression was increased due to autophagy block (**Figures 4G and H**). Moreover, puromycin-labeling of Plin2 showed was increased during NASH (**Figures 4I**), confirming their regulation by other translation factors such as eIF6, eIF2, RPS6KB1 [17, 24, 26].

**Fig. 4.**
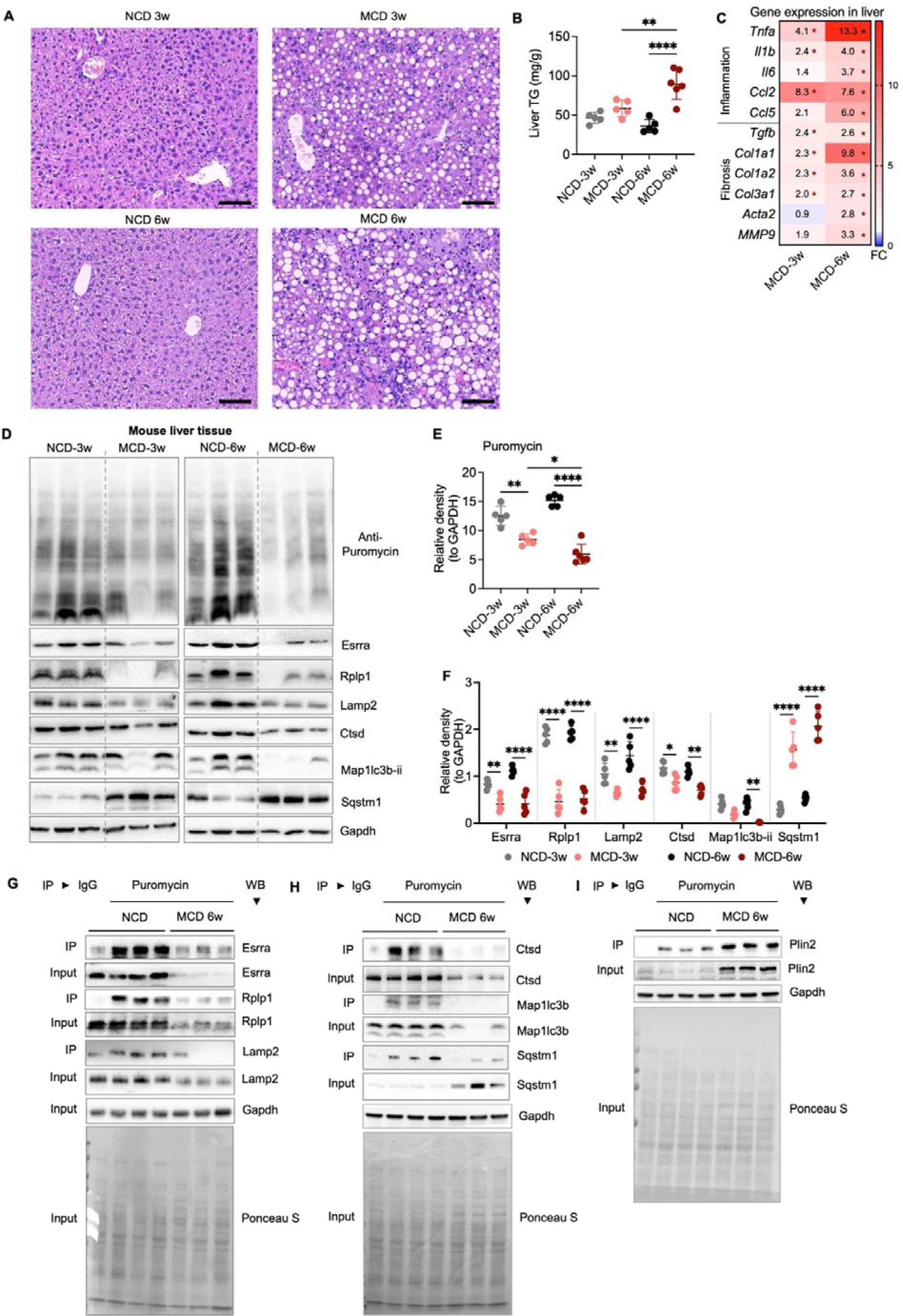
Defective Esrra-Rplp1 pathway leading to decreased translation rate and lysosome-autophagy proteins in MCD diet fed mice livers. (A) Representative micrograph of H&E staining in liver tissues collected from mice fed NCD or methionine & choline-deficient diet (MCD) diet for 3 and 6 weeks (n=5 per group). Scale bars are 100 μM. (B) Triglycerides (TG) measurement in the Liver tissues. (C) Heat map representing RT-qPCR analysis of inflammatory and fibrosis genes in the liver tissues from mice fed MCD-diet for 3 or 6 weeks (n=5 per group). Gene expression is normalized to Gapdh. (D) Representative Western blots of livers from mice fed NCD or MCD diet for 3 and 6 weeks (n=5 per group). (E and F) Plots represent relative density of corresponding Western blots normalized to Gapdh. (G, H and I) Western blots of immunoprecipitated (IP) proteins labelled by puromycin or IgG. Puromycin labeling shows recent translation events. Levels of significance: *P<0.05; **P<0.01; ***P<0.001; ****P<0.0001.

### 2.5 Esrra regulated Rplp1-dependent translation of lysosome/autophagy proteins in mice model of NASH

To understand Esrra’s role in regulating Rplp1 and lysosome/autophagy proteins’ expression *in vivo*, first we generated liver-specific knock out mice. Mice with liver-specific deletion of *Esrra* (LKO) had increased liver index and liver triglycerides (TG), and decreased β-hydroxybutyrate when fed normal chow diet (NCD; age 20-22 weeks) and compared to their wild-type (WT) littermates (**Supplementary Figures 5A-C**). This indicated that LKO had fatty liver and decreased β-oxidation of fatty acids even on NCD. Interestingly, LKO mice also had an increased inflammatory and fibrosis gene expression (**Supplementary Figure 5D**). Furthermore, these LKO mice had decreased protein expression of Rplp1, Lamp2, Ctsd and Map1lc3b-ii protein and accumulation of Sqstm1 suggesting a late block in autophagy due to impaired lysosomal function (**Supplementary Figures 5E and F**). However, mRNA expression of only Rplp1 was significantly decreased whereas mRNA expression of Lamp2, Ctsd, Map1lc3bii, and Sqstm1 expression was not changed significantly (**Supplementary Figure 5G**). These data confirm that Esrra transcriptionally regulated *Rplp1*, whereas the lysosome and autophagy protein expression were regulated at translational level. Further, the data also suggested that basal autophagic impairment in the LKO mice could increase hepatic inflammation and fibrosis as previously reported [2, 10, 11, 15, 21].

We next examined liver-specific *(Alb)-Esrra* overexpression in mice fed WDF and found that it significantly increased hepatic Rplp1, Lamp2, Ctsd, and Map1lc3b-ii protein expression compared to control null-mice fed WDF (**Figures 5A and B**). The Sqstm1 protein accumulation found in null-mice fed WDF decreased in *(Alb)-Esrra* overexpressing mice fed WDF in conjunction with the increase in Map1lc3b-ii expression, and suggested there was improved autophagy flux in these mice. H&E staining showed improvements in NASH pathological features such as micro/macro steatosis, fatty hepatocytes hypertrophy/ballooning and inflammatory foci in *(Alb)-Esrra* overexpressing mice compared to null-mice when both were fed WDF (**Figure 5C).** Furthermore, *(Alb)-Esrra*-overexpressing mice fed WDF had reduced liver index, liver TG, and serum glucose and increased serum β-HB compared to control mice when both were fed WDF (**Figures 5D-G**). *(Alb)-Esrra*-overexpressing mice fed WDF had reduced hepatic inflammation and fibrosis gene expression compared to null-mice when fed WDF suggesting that Esrra overexpression improved NASH *in vivo* (**Figure 5H**). Previously, Esrra inhibition showed improvement during hepatosteatosis in a global *Esrra*-KO mouse model [48] fed high fat diet (HFD), whereas pharmacological inhibition worsened rapamycin-induced fatty liver [37]. Of note, we examined liver-specific inactivation/activation of Esrra rather than systemic inhibitory effects of Esrra on NASH.

**Fig. 5.**
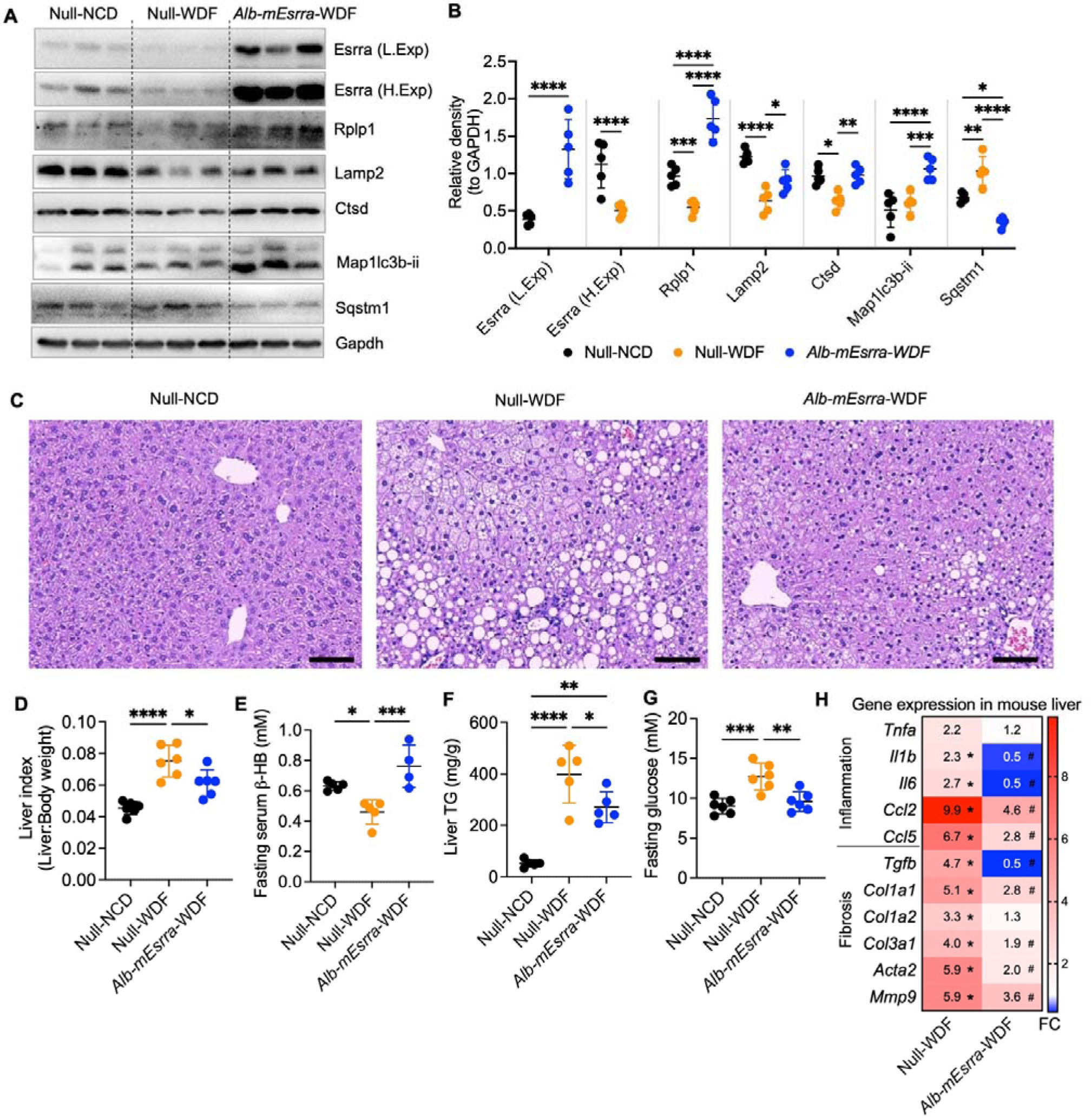
Hepatic *Esrra* genetic deletion or overexpression positively regulated Rplp1, lysosome, and autophagy proteins’ expression *in vivo*. (A) Representative Western blots of livers from null and hepatic-Esrra overexpressed mice fed NCD or WDF for 16 weeks (n=5 per group). Esrra Western blots were low exposed (L. Exp) and high exposed (H. Exp) for accurate densitometric measurements of Esrra overexpression and basal Esrra expression. (B) Plots represent relative density of corresponding Western blots normalized to Gapdh. (C) Representative micrograph of H&E staining in liver tissues collected from null and hepatic-Esrra overexpressed mice fed NCD or WDF for 16 weeks (n=5 per group). Scale bars are 100 μM. (D-G) Measurements of liver index, serum β-HB, liver TG, and fasting blood glucose measurements in null and hepatic-Esrra overexpressed mice fed NCD or WDF for 16 weeks.(H) RT-qPCR analysis of relative gene expression in livers from null and hepatic-Esrra overexpressed mice fed NCD or WDF for 16 weeks (n=5 per group). Gene expression was normalized to *Gapdh*. Levels of significance: *P<0.05; **P<0.01; ***P<0.001; ****P<0.0001.

### 2.6 Alternate day fasting-improved NASH required activation of Esrra/Rplp1-mediated translation

Intermittent fasting regimens involving alternate day fasting or time-restricted feeding for several days have had beneficial effects in obesity, diabetes, and NASH [49–51]. Moreover, these beneficial effects induced by fasting have been associated with activation of autophagy [16, 49, 52–55]. Esrra protein is induced by fasting/starvation (**Figure 6A**) [48]; and, it is noteworthy that hepatic *Esrra* overexpression in mice fed WDF activated the Esrra-Rplp1-lysosome pathway, improved autophagy, and reduced hepatosteatosis, inflammation, and fibrosis compared to null mice when both were fed WDF (**Figure 5A-H**). Thus, we examined whether intermittent fasting could induce the Esrra-Rplp1-lysosome pathway and have beneficial effects in mice with a pre-established NASH condition. Accordingly, we employed an alternate day fasting regimen for five cycles in mice earlier fed WDF for 16 weeks to pre-establish NASH. These mice with NASH then were injected with vehicle or Esrra-specific inhibitor C29 during fasting to examine the role of Esrra during intermittent fasting (**Figure 6B**). Mice fed NCD or WDF continuously were used as controls. Mice fed WDF for 16 weeks developed NASH and had significantly reduced puromycin-labeling of overall proteins and the decreased protein expression of Esrra, Rplp1, Lamp2, Ctsd and Maplc3b-ii suggesting there was impaired lysosome-autophagy protein translation rates and function at baseline (**Figures 6C-F**). Surprisingly, alternate day fasting for five cycles induced the expression of Esrra and Rplp1, and reversed the decline in puromycin-labeled proteins in mice fed WDF. This regimen also restored Rplp1, Lamp2, Ctsd, Map1lc3b-ii proteins’ expression and decreased Sqstm1 protein expression suggesting activation of lysosome and improved autophagy (**Figures 6E and F**). In contrast, injection of the Esrra inhibitor, C29, during only fasting (when Esrra expression was increased) significantly inhibited Esrra protein expression and increased Sqstm1 expression suggesting there was impaired autophagy. We further confirmed that the protein expression in C29 injected mice was attenuated at translation level as the gene expression of Lamp2, Ctsd, Map1c3b and sqstm1 were still upregulated in C29 treated mice (**Supplementary Figure 6**).

**Fig. 6.**
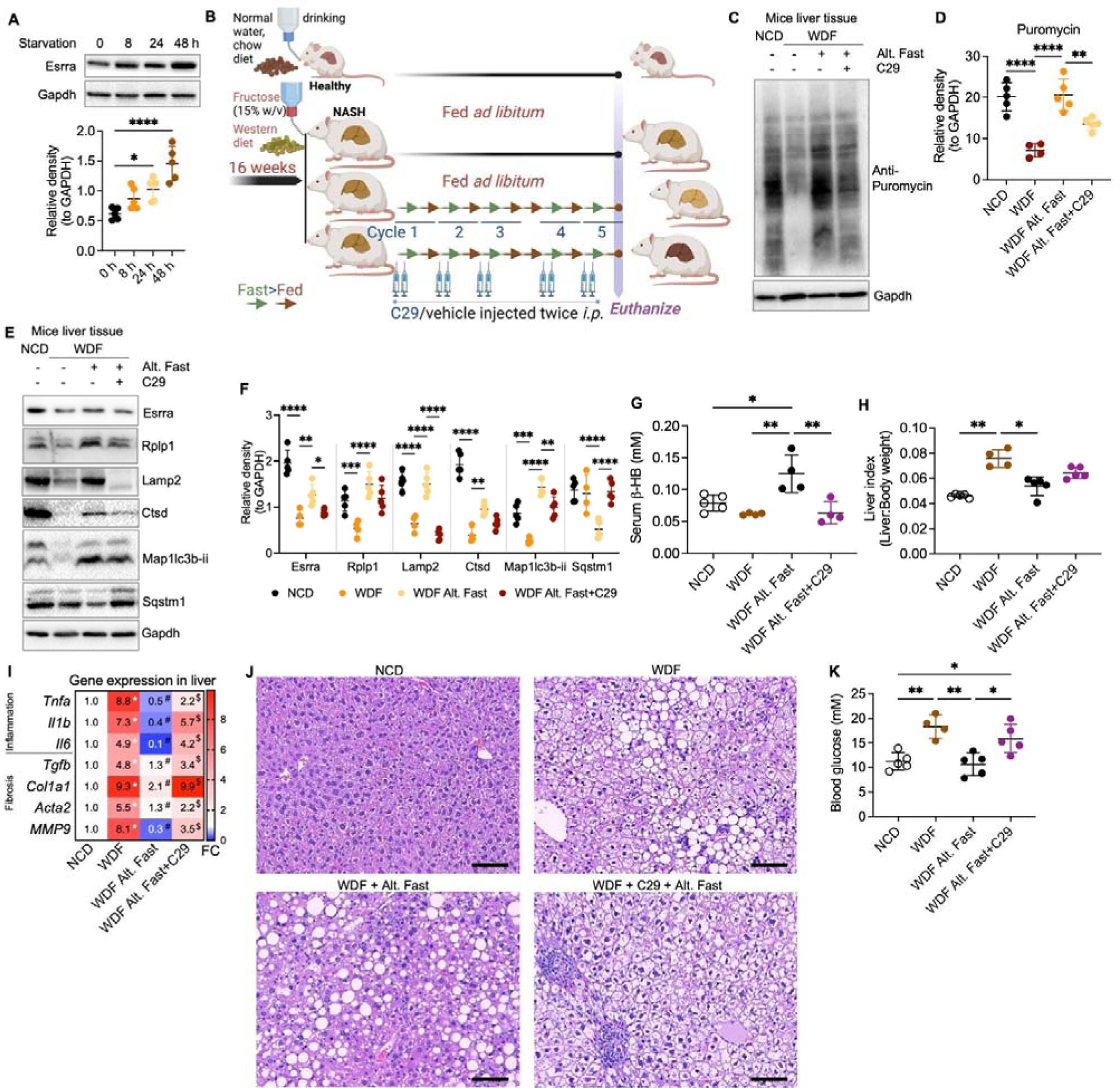
Esrra-Rplp1 axis was required for improved translation and lysosome-autophagy during alternate day fasting in NASH liver. (A) Representative Western blots of livers from starved mice for indicated time points. Dot plots represent relative density of Western blot normalized to Gapdh (n=5 per group). (B) Scheme shows alternate day fasting regime in NASH mice: WDF fed mice for 16 weeks fasted and fed alternate days and injected vehicle or C29 (to inhibit Esrra activity) during fasting days only. After five Fast-Fed cycles, blood and liver tissue were harvested for analysis. Untreated mice fed NCD or WDF for 16 weeks were used for comparison (n=4-5 per group). (C) Representative Western blots of livers from puromycin injected WDF fed mice (n=4-5 per group) fasted and fed alternate days for 5 consecutive cycles and injected vehicle or C29 during fasting days (as shown in Panel A). Untreated mice fed NCD or WDF for 16 weeks were used for comparison (n=4-5 per group). (D) Plot represents relative density of Western blot normalized to Gapdh. (E) Representative Western blots of livers as described in panel A. (F) Plots represent relative density of Western blots normalized to Gapdh. (G and H) Serum β-HB measurement and liver index measurements in the livers from the mice described in panel B. (I) RT-qPCR analysis of relative gene expression in the livers from the mice described in panel A. Gene expression was normalized to *Gapdh* (n=4-5 per group). (J) Representative (n=4-5 per group) micrograph of H&E staining in liver tissues collected from mice as described in panel A. Scale bars are 100 μM. (K) Blood glucose measurements in the livers from the mice described in panel B. Levels of significance: *P<0.05; **P<0.01; ***P<0.001; ****P<0.0001.

Remarkably, mice fed WDF that underwent alternate day fasting had significant improvements in serum β-HB, liver index, and serum glucose levels in conjunction with the increased Esrra protein expression induced by alternate day fasting **(Figures 6G, H, and K)**. In contrast, mice fed WDF that underwent alternate-day fasting and C29 injection during alternate day fasting had little or no improvement in these parameters compared to mice fed WDF continuously. Additionally, mice fed WDF undergoing alternate day fasting while fed WDF had reduced expression of inflammatory and fibrosis genes compared to mice fed WDF continuously or mice fed WDF and injected with C29 while undergoing alternate day fasting (**Figures 6I)**. H&E staining also showed that alternate day fasting improved NASH histological features (micro/macro steatosis and inflammatory foci) in WDF fed mice, while C29 treatment worsened these features (**Figures 6J).** Thus, alternate day fasting induced Esrra expression and restored Rplp1-mediated translation of lysosome-autophagy proteins and decreased inflammation and fibrosis. Taken together, we now have identified a novel pathway that plays a key role in the beneficial fasting-mediated improvements on metabolism and NASH. It is noteworthy that the induction of Esrra and the inhibition of mTOR activity also may have synergistic effects during fasting [37, 48].

## 3. Conclusion

Our investigation into the pathophysiology of NASH has unveiled a critical role for the nuclear hormone receptor Esrra in controlling key cellular mechanisms that maintain liver health. We showed that there was decreased lysosome/autophagy proteins translation in NASH that was dependent upon the induction of Rplp1 expression by Esrra (**Supplementary Figure 7**). This deficit hampers autophagy and β-oxidation of fatty acids, processes essential for the prevention of fat accumulation, inflammation, and fibrosis in the liver-characteristic features of NASH [1]. Intriguingly, our results indicate that the adverse effects on protein translation seen in NASH can be reversed. Overexpression of Esrra in cellular and mouse models restore translation defects and reinstate normal autophagy and lipid metabolism pathways, highlighting a potential therapeutic target. Other important finding is that alternate day fasting promotes Esrra and Rplp1 expression, which in turn restores normal protein expression, thus offering a dietary approach to manage NASH. The mechanism(s) for the beneficial effects of intermittent fasting on NASH are poorly understood but likely involve the induction of the Essra-Rplp1-lysosome pathway to restore autophagy and β-oxidation of fatty acids.

Nuclear receptors have traditionally been classified as transcription factors that are activated by specific ligands [56]. Yet, our research reveals a multifaceted role for the nuclear receptor Esrra, showing that it governs not only gene transcription but also plays a pivotal role in the translation of crucial proteins. This regulatory effect on protein synthesis occurs through an intermediary mechanism that necessitates the activation of Rplp1. Our study proposes that agents capable of stimulating Esrra expression or functioning as Esrra agonists could potentially initiate the Esrra-Rplp1-depnedent translation of lysosome/autophagy proteins, thereby offering a novel approach to slow or potentially reverse the progression of NASH. This could be especially effective when used in conjunction with dietary modifications, such as alternate day fasting.

Previously, we have demonstrated that thyroid hormone (TH) enhances the expression of Ppargc1a and Esrra [23], thereby not only activating hepatic autophagy and mitochondrial function but also potentially engaging both protein translation and gene transcription pathways that are instrumental in ameliorating NASH [21, 57, 58]. Whether the disruption of the Esrra-Rplp1 pathway is a disruption exclusive to NASH or if it is implicated in other liver-related, metabolic, or cancerous conditions remains unknown. Esrra’s role in inducing autophagy across a range of cell types [23, 34–36, 59, 60] invites further investigation into its broader biological implications.

This research offers significant clinical implications in light of the current absence of pharmacological options for NASH. By demonstrating that Esrra is a dual regulator-impacting both transcription and translation mechanisms-it emphasizes Esrra’s central role in maintaining hepatic cellular homeostasis. Our findings not only underscore the critical involvement of Esrra in the development of NASH but also suggest new therapeutic possibilities that leverage the body’s intrinsic response to fasting. Moreover, they prompt further exploration into whether the Esrra-Rplp1 axis plays a universal role in various diseases, potentially transforming our strategies for treating a broad range of metabolic disorders.

## 4. Methods

### 4.1 Animal studies

#### 4.1.1 MCD diet

Mice were fed with normal chow control diet (NCD) or methionine- and choline-deficient L-amino acid diet (MCD; A02082002BR, Research Diets, Inc.) for 3 and 6 weeks for progressively develop NASH [39, 61].

#### 4.1.2 WDF diet

Mice were fed with normal chow control diet (NCD) or Western diet (WD; D12079B, Research Diets, Inc.) supplemented with 15% (w/v) fructose (F0127, Sigma-Aldrich) in drinking water (WDF) for 8 or 16 weeks to progressively generate steatosis and NASH respectively [6, 7, 61].

#### 4.1.3 Starvation and alternate day fasting study

10-12 weeks old male mice were fasted overnight for food synchronization two nights before starvation for 0, 8, 24, and 48 h. For alternate day fasting, mice were fed WDF for 16 weeks, and fasted and fed alternate days (Fast-Fed one cycle) for five cycle (Figure 6B). We injected C29 (10 mg/kg body weight) or vehicle *i.p.* twice a day. Puromycin (Puro, 20 mg/kg body weight) was injected *i.p.* 30 min before euthanization for protein-labeling and translation analysis. Mice were euthanized during fed condition.

#### 4.1.4 Liver-specific <u>Esrra</u> <u>KO mice (LKO)</u>

*Esrra* WT and LKO mice are described in the supplementary data, were fed normal chow diet and fasted 6 h before euthanization.

#### 4.1.5 Alb-mESRRA overexpression

Mice were generated by injecting AAV8-*Alb*-*mEsrra* (5X1011 gc/mice) via tail vein and housed for four weeks with no other intervention [6].

#### 4.1.6 General mouse care and ethics statement

Mice were simple randomized before grouping and fed different diets and normal water, or fructose treated *ad libitum*. All mice were maintained according to the Guide for the Care and Use of Laboratory Animals (NIH publication no. One.0.0. Revised 2011), and the experiments performed were approved by the IACUCs at SingHealth (2015/SHS/1104) and (2020/SHS/1549). Other details can be found in Supplementary Methods.

### 4.2 Cell cultures

AML12 cells (ATCC® CRL-2254™) and Primary human hepatocytes (5200, ScienCell) were cultured as described in the Supplementary Methods.

4.3 Other methodological details for Label-Free Quantification (LFQ) of hepatic proteome, Polysome profiling, ChIPseq, ChIP-qPCR analysis, *in vitro* gene manipulation, measurements of blood glucose, serum β-hydroxybutyrate (β-HB/Ketone bodies), and serum and liver triglycerides (TG), RNA/protein expression, statistical analysis etc. can be found in the Supplementary Methods section of Supplementary Information file.

## Acknowledgements and funding details

The authors like to acknowledge that the research is funded by the Ministry of Health (MOH), and National Medical Research Council (NMRC), Singapore, grant number NMRC/OFYIRG/0002/2016 and MOH-000319 (MOH-OFIRG19may-0002), Duke/Duke-NUS Research Collaboration Pilot Project Award (Duke/Duke-NUS/RECA(Pilot)/2022/0060), and KBrFA (Duke-NUS-KBrFA/2023/0075) to BKS; NMRC/OFYIRG/077/2018 to MT; and CSAI19may-0002 to PMY; Duke-NUS Medical School and Estate of Tan Sri Khoo Teck Puat Khoo Pilot Award (Collaborative) Duke-NUS-KP(Coll)/2018/0007A to JZ. This work is also partially supported by grants from the Louisiana Clinical and Translational Science Center (NIGMS 2U54GM104940), the National Heart Lung and Blood Insititute, NIH, USA (NHLBI R01HL146462-01), and the Khoo Bridge Fund, Singapore (KBrFA/2022/0060) to SG. The illustrations were made on BioRender.com.

## Author’s contributions

MT, BKS, LW, SG, experimental design/execution, data analysis; KG, Esrra KO mice experiments; KT, RS, KA, wet lab work; JZ, hepatic-ESRRA overexpressing mice; WWT, YTN, LS, polysome profiling; VG, generated ESRRA KO mice, finalized manuscript; SP, DPM, shared C29, experimental suggestions, finalized manuscript; YW, BHB, EM, finalized manuscript; MT, BKS, PMY, drafted/finalized manuscript, provided financial support.

## Disclosure

Authors have no conflict of interests.

## Notes

### Competing Interest Statement

The authors have declared no competing interest.

### Summary of Updates

In accordance with the valuable feedback provided by the reviewers, we have revised the manuscript to exclusively focus on the NASH studies, separating out the aspects related to starvation studies for enhanced clarity and coherence. The findings from the starvation studies are being prepared for submission in a distinct manuscript to ensure a more focused and comprehensive understanding of each subject. Additionally, we have supplemented our submission with newly generated in vivo data on NASH, further enriching the content and integrity of this standalone NASH-focused manuscript.

