## Supplemental data and methods for "Esrra regulates Rplp1-mediated translation of lysosome proteins suppressed in non-alcoholic steatohepatitis and reversed by alternate day fasting"

*^1^Cardiovascular and Metabolic Disorders Program, Duke-National University of Singapore (NUS) Medical School, Singapore 169857, Singapore. ^2^Institut de Génomique Fonctionnelle de Lyon, Université de Lyon, Université Lyon 1, CNRS, Ecole Normale Supérieure de Lyon, 46 Allée d’Italie, 69364 Lyon Cedex 07, France. ^3^Department of Pharmacology and Cancer Biology, Duke University School of Medicine, C238A Levine Science Research Center, Durham, NC 27710, USA. ^4^Department of Anatomy, Yong Loo Lin School of Medicine, NUS, Singapore 117594. ^5^Centre for Computational Biology, Cardiovascular and Metabolic Disorders Program, Duke–National University of Singapore (NUS) Medical School, Singapore 169857, Singapore. ^6^Goodman Cancer Research Centre, McGill University, 1160 Pine Avenue West, Montreal, Québec H3A 1A3, Canada. ^7^Dept of Surgery, Singapore General Hospital and Dept. of Surgical Oncology, National Cancer Centre, Singapore-169608*,*^8^Duke Molecular Physiology Institute and Dept. of Medicine, Duke University School of Medicine, Durham, NC 27710, USA.*

**Supplementary material contains:**

**Supplementary Figures (1, 2, 3, 4, 5 and 6)**

**Supplementary methods**

**Supplementary Tables (1 and 2)**

**Supplementary Figures**

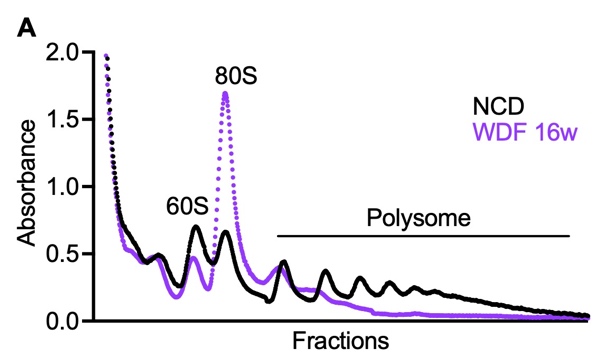

**Supplementary Fig. 1. Global protein translation was decreased in the livers of NASH mice fed WDF for 16 weeks.**  (A) Representative polysome profile of the liver tissues collected from mice fed NCD or WDF for 16 weeks.

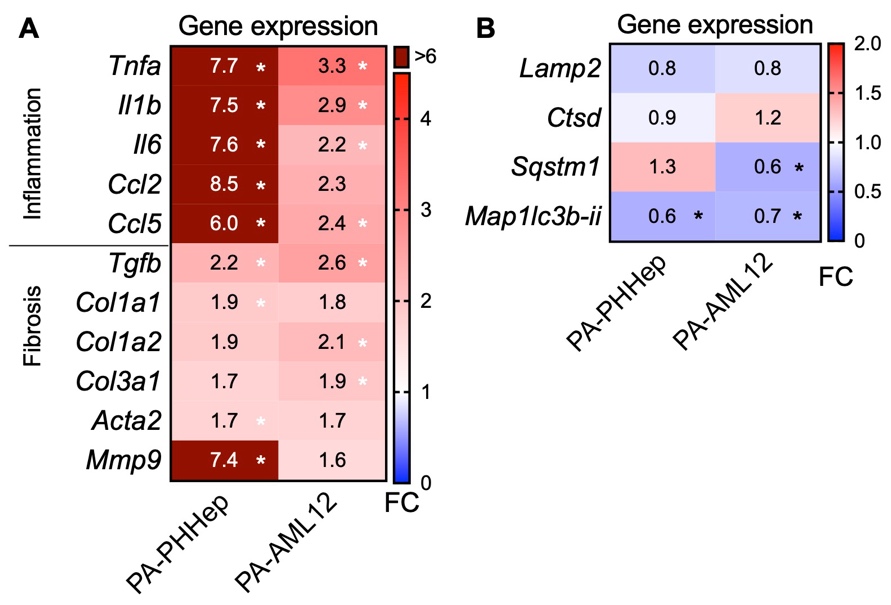

**Supplementary Fig. 2. Protein translation was decreased in lipotoxicity and regulated by Esrra-Rplp1 axis.**  (A and B) RT-qPCR analysis of relative gene expression in 24 h PA (0.5 mM) treated primary human hepatocytes (PHHep; n=3-5 per group) and AML12 cells (n=3-5 per group). Gene expression is normalized to Gapdh.

Levels of significance: *P<0.05; *Compared to the control.

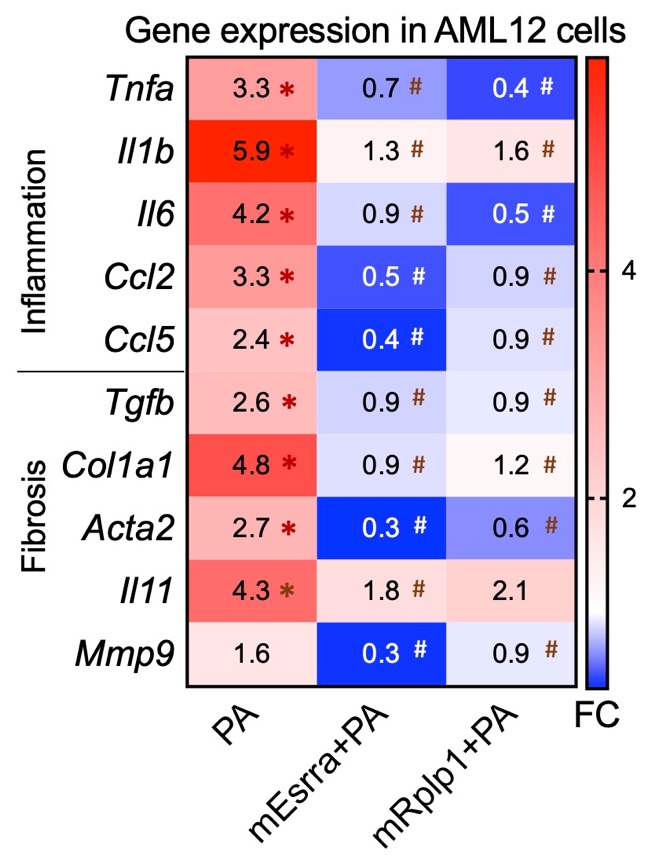

**Suppl. Fig. 3. Esrra or Rplp1 overexpression increased translation and lysosome-autophagy acticity in hepatic cells.** (A) RT-qPCR analysis of relative gene expression in 24 h PA (0.5 mM) treated AML12 cells with overexpressions of Esrra or Rplp1 or empty vector.

Levels of significance: *P<0.05; **P<0.01; ***P<0.001; ****P<0.0001, *Compared to the control.

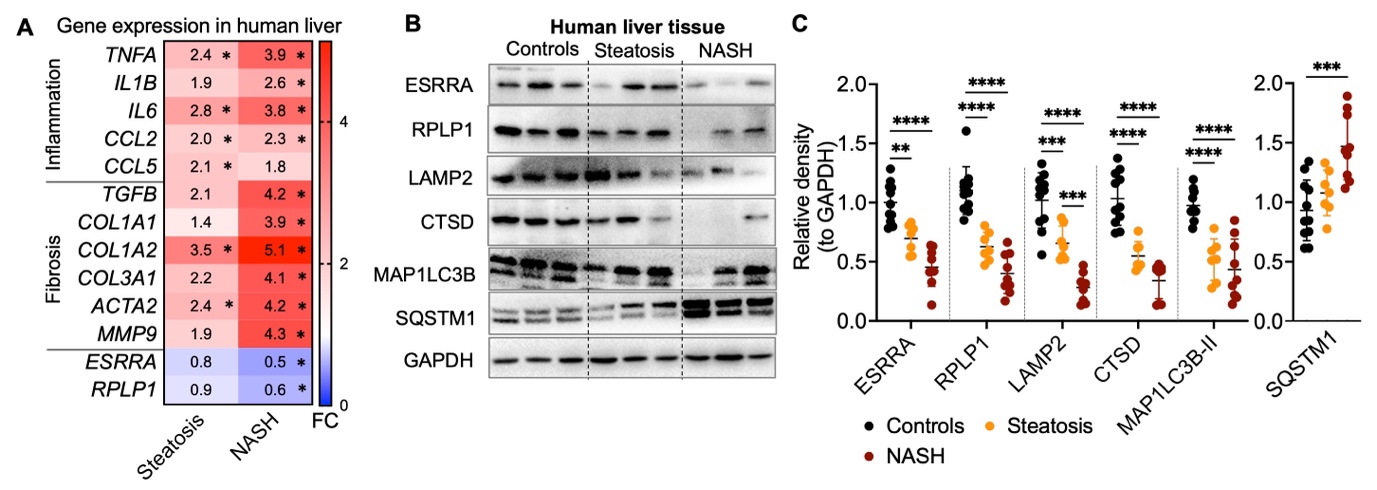

**Suppl. Fig. 4. Defective Esrra-Rplp1 pathway leading to decreased translation of lysosome-autophagy proteins in the livers of NASH patients.** (A) RT-qPCR analysis of relative gene expression in human livers collected from steatosis (n=7) and NASH (n=7) patients compared to control (n=6) subjects. Gene expression was normalized to *POLR2A*. (B) Representative Western blot of human livers collected from steatosis (n=7) and NASH (n=9) patients compared to control (n=11) subjects. (C) Plots represent relative density of corresponding Western blots normalized to Gapdh.

Levels of significance: *P<0.05; **P<0.01; ***P<0.001; ****P<0.0001, *Compared to the control.

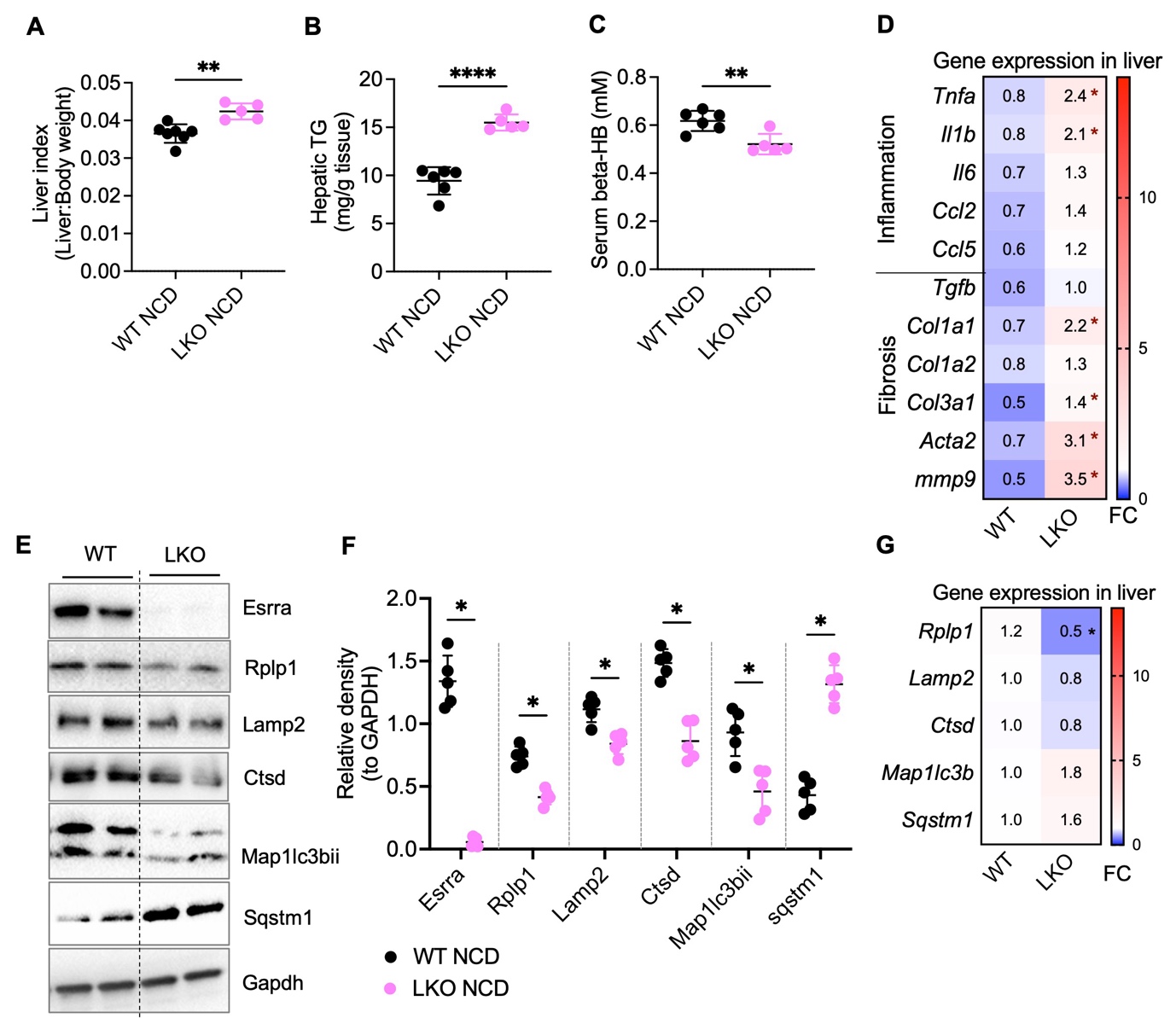

**Suppl. Fig. 5. Hepatic Esrra deletion led to increased inflammatory and fibrosis genes with decreased translation of lysosome-autophagy proteins in liver.** (A and B) Liver index (liver weight:body weight) of wild type (WT) and liver-specific *Esrra* knockout (LKO) mice fed normal chow diet (NCD) (n=7-5 per group). (B and C) Measurements of hepatic triglycerides (TG) and serum β-hydroxybutyrate (β-HB) from WT and LKO mice (n=6-5 per group). (D) RT-qPCR analysis of relative gene expression in livers collected from hepatic Esrra knockout mice (n=5 per group). (E) Representative Western blots of livers collected from WT and LKO mice (n=5 per group). (F) Plots represent relative density of corresponding Western blots normalized to Gapdh. (G) RT-qPCR analysis of relative gene expression in livers collected from hepatic Esrra knockout mice (n=5 per group). Gene expression was normalized to *Polr2a*.

Levels of significance: *P<0.05, *Compared to the control.

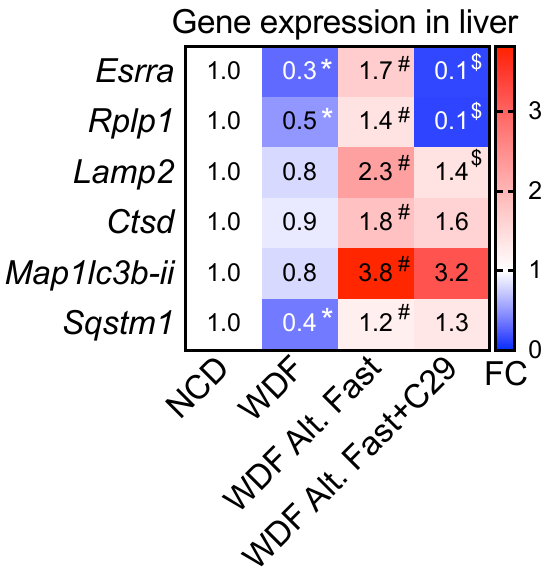

**Suppl. Fig. 6. Hepatic Esrra deletion led to increased inflammatory and fibrosis genes with decreased translation of lysosome-autophagy proteins in liver.** RT-qPCR analysis of relative gene expression in livers collected from the mice showed in the main Figure 6 (panel B), (n=4-5 per group). Gene expression was normalized to *POLR2A*.

Levels of significance: *P<0.05, *Compared to the control.

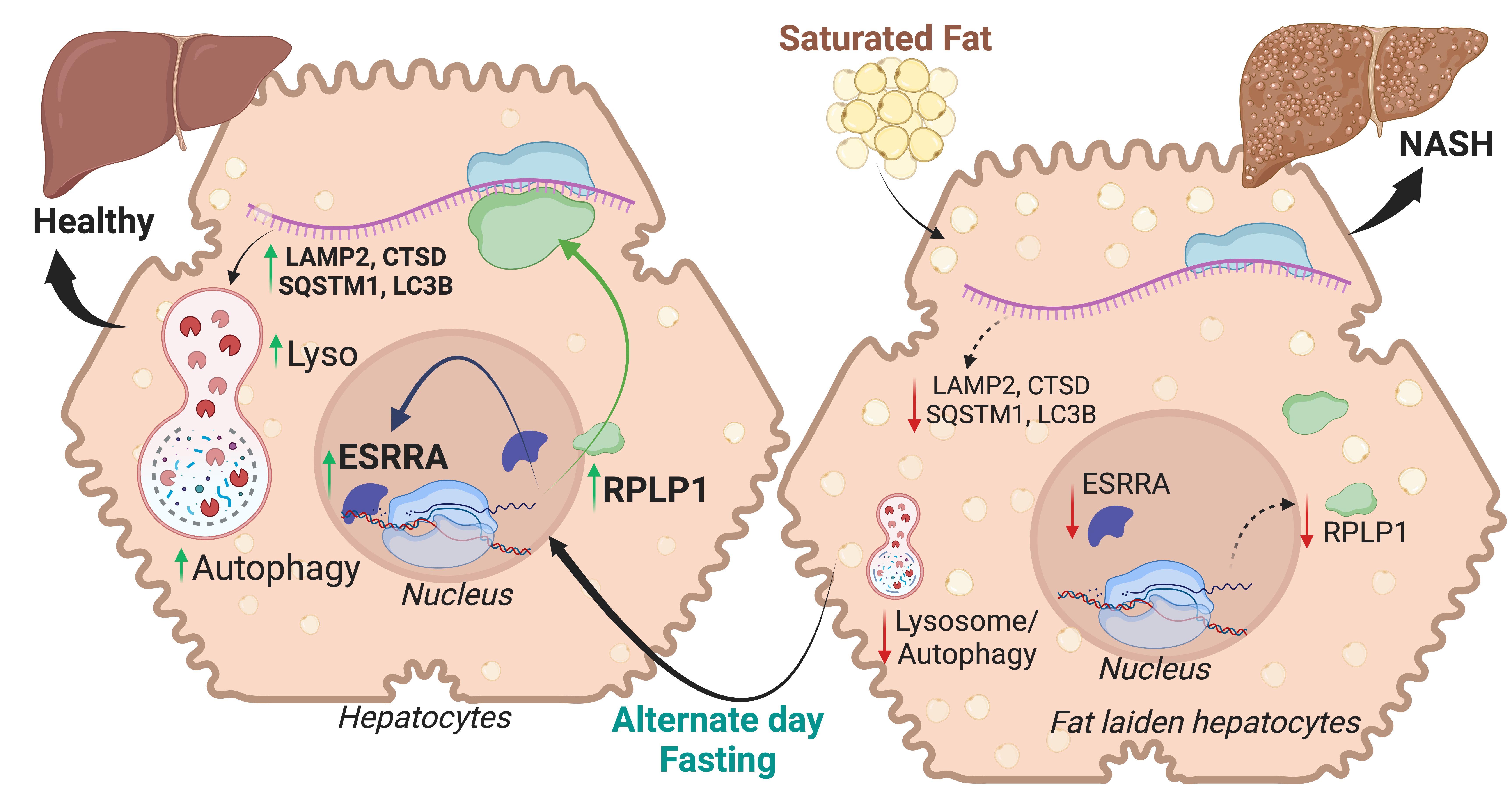

**Suppl. Fig. 7. Illustration summarizing Esrra-Rplp1-lysosome axis regulation under alternate day fasted or NASH conditions.** Our findings show Esrra activation is required for later upregulation of ribosomal proteins including Rplp1 during alternate day fasting that leads to transcriptional activation of lysosomal and autophagy proteins. Translational recovery of lysosomal and autophagy proteins is essential for metabolic homeostasis. However, in pathological conditions such as NASH Esrra/Rplp1-mediated translation is impaired leading to lower lysosomal-autophagy proteins causing lipid accumulation, inflammation, fibrosis and cellular damage in NASH progression. Esrra activation by alternate day fasting in NASH restores Esrra-Rplp1-Lysosome-Autophagy pathway and result in decreased hepatic steatosis, inflammation, fibrosis, thus NASH regression towards healthy liver phenotype.

**Supplementary Methods**

*Animal studies*

MCD diet: 8-10 weeks old male C57BL/6J mice (n-5/group) were fed with normal chow control diet (NCD) or methionine-and choline-deficient L-amino acid diet (MCD; A02082002BR, Research Diets, Inc.) for 3 and 6 weeks for progressively develop NASH [1, 2]. Puromycin (Puro, 20 mg/kg body weight) was injected *i.p.* 30 min before euthanization for rate of translation analysis and cycloheximide (CHX, 0.1 mg/g body weight) was injected *i.p.* 60 min before euthanization to inhibit translation [3, 4]. Later, mice were fasted for 6h and euthanized, blood and liver tissues were collected for mRNA and protein analysis.

WDF diet: 8-10 weeks old male C57BL/6J mice (n-5/group) were fed with normal chow control diet (NCD) or Western diet (WD; D12079B, Research Diets, Inc.) supplemented with 15% (w/v) fructose (F0127, Sigma-Aldrich) in drinking water (WDF) for 8 or 16 weeks to progressively generate steatosis and NASH respectively [2, 5, 6]. Later, mice were fasted for 6h and euthanized, blood and liver tissues were collected for mRNA and protein analysis.

Starvation study: 10-12 weeks old male mice were fasted overnight for food synchronization and fed to two days *ad libitum*. Mice were starved for 0, 8, 24, and 48 h (n=5 per group) starting from 10:00AM and euthanized at 6:00PM same day, 10:00AM next day and a day after respectively.

Alternate day fasting study: Mice were fed WDF for 16 weeks to induce NASH or NCD as control. After 16 weeks, mice were fasted and fed alternate days (Fast-Fed one cycle) for five cycle (n=4-5 per group). For inhibiting Esrra specifically during fasting, we injected C29 (10 mg/kg body weight) or vehicle *i.p.* twice a day (5 h before the start and just before the start of each fasting cycle) in only fasting groups during fasting day while there was no injection given during fed days. Puromycin (Puro, 20 mg/kg body weight) was injected *i.p.* 30 min before euthanization for protein-labeling and translation analysis. Mice were euthanized during fed condition. Mice were euthanized during fed condition, blood and liver tissues were collected for mRNA and protein analysis.

Liver-specific Esrra knockout (KO) mice generation: We obtained Esrra flox/flox mice from Dr. Jean-Marc Vanacker (Université de Lyon, France.) and described elsewhere [7]. We crossed these Esrra flox/flox mice with Alfp-Cre transgene where the Cre recombinase was expressed under the control of the mouse *albumin* enhancer and promoter and the mouse *alpha-fetoprotein* enhancers as decribed earlier [8]. We used ERRA Flox/Flox mice as the wild type control and Esrra^Fl/Fl^:Alfpcre^+^ (heterozygote) as the liver-specific LKOs.

Esrra WT and Esrra LKO mice (n-5/group) were fed normal chow diets 18-20 week of age, fasting overnight and euthanized, blood and liver tissues were collected for mRNA and protein analysis.

*Alb*-mESRRA overexpression: For liver-specific expression, we used AAV8 mediated gene delivery of m*Esrra* gene cloned under the control of mouse *Alb* promoter (Vector Biolabs, USA). 8 weeks old male C57BL/6J mice (n-5/group) were used for liver-specific *Esrra* overexpression (*Alb-mEsrra*). Mice were generated by injecting AAV8-*Alb*-*mEsrra* (5X1011 gc/mice) via tail vein and housed for four weeks with no other intervention [5]. We injected AAV8-*Alb*-Null which does not contain any DNA sequence under the transcriptional control of the *Alb* promoter as controls. Later, mice were euthanized, blood and liver tissues were collected for mRNA and protein analysis.

General mouse care and ethics statement: mice were purchased from InVivos, Singapore and, housed in hanging polycarbonate cages under a 12 h/12 h light/dark schedule at Duke-NUS vivarium. Mice were simple randomized before grouping and fed different diets and normal water, or fructose treated *ad libitum*. Animals were euthanized in CO2 chambers. All mice were maintained according to the Guide for the Care and Use of Laboratory Animals (NIH publication no. One.0.0. Revised 2011), and the experiments performed were approved by the IACUCs at SingHealth (2015/SHS/1104) and (2020/SHS/1549).

*Blood glucose, serum β-hydroxybutyrate (β-HB/Ketone bodies), and serum and liver triglycerides (TG) measurements.*

Blood glucose was measured from euthanized mice during the blood collection using Roche’s strip-based glucometer (Accu-Chek Performa, Roche). Serum β-HB and TG were measured using β-HB (Ketone Body) Colorimetric Assay Kit (#700190, Cayman) and Triglyceride Colorimetric Assay Kit (#10010303, Cayman).

*Cell cultures*

AML12 cells (ATCC® CRL-2254™) were cultured as indicated elsewhere [9, 10]. Starvation medium (DMEM:F12 mix with Pen/Strep lacking serum, ITS and Dexamethasone) was used for serum starvation experiments. To analyze the rate of translation, puromycin (10 μg/ml) was added for 15 min before harvest [11] whereas C29 (5 μM) was added to media for indicated time to inhibit Esrra [12]. Palmitic acid (PA) 0.5-0.75 mM conjugated with 0.5% BSA was used for 24 h as described elsewhere [5]. 0.5% BSA solution was used as control (CT). Bafilomycin A1 (5 nM) was used to analyze autophagy flux [13].

Primary human hepatocytes (5200, ScienCell) were cultured as indicated elsewhere [5, 14]. Puromycin (10 μg/ml) was added for 10 min before harvest [11] whereas C29 (5 μM) was added to media for indicated time [12]. PA (0.5mM) was used as stated above.

*Gene manipulation in cultured cells*

Gene knockdown in vitro: Silencer Select siRNAs (s4829, s4830, and s4831; Life Technologies Inc.) or ON-TARGETplus Smartpool siRNAs (L-040772-00-0010, Dharmacon) against *Esrra*, and Silencer Select siRNA (s234520; Life Technologies Inc.) for *Rplp1* gene knockdown were used in AML12 cells. Negative siRNA (Silencer Negative Control No. 1 siRNA; AM4611, Life Technologies Inc.) was used as a negative control. Transfections were carried out in AML12 cells in a 12-well or 6-well plate or four-well chambered slides or 24-well Seahorse XF plate using 30 nM of the above indicated siRNAs and negative control siRNA with Lipofectamine RNAiMAX (Invitrogen; Life Technologies Inc.) following the reverse transfection protocol, as indicated elsewhere [9, 13].

Gene overexpression *in vitro*: ORF sequence of m*Esrra* (Gene ID: 26379) and m*Rplp1* (Gene ID: 56040) genes were cloned in a pcDNA3.1(+)-C-6His plasmid by Genescript Limited (Hong Kong) to get Esrra_OMu13026C_pcDNA3.1(+)-C-6His and Rplp1_OMu11118C_pcDNA3.1(+)-C-6His constructs for *Esrra* and *Rplp1* overexpression in AML12 cells respectively. Transfections were carried out in a 12-well plate or 6-well plate using Lipofectamine 3000 (Invitrogen; Life Technologies Inc.) following the reverse transfection protocol, as described elsewhere [13]. Empty pcDNA3.1(+)-C-6His vector was used as control.

*RNA isolation and RT-qPCR analysis of gene expression*

RNA isolation and RT-qPCR for measuring gene expression were performed as described earlier [13]. Predesigned KiCqStart SYBR Green optimized primers from Sigma-Aldrich (KSPQ12012) were used for RT-qPCR.

*Protein isolation and Western blotting analysis of protein expression*

Protein isolation and Western blotting analysis of protein expression were performed as described earlier [13]. Primary antibodies for puromycin (1:25,000 dilution; MABE343, Merck), 1:1000 dilution of Esrra (ab16363, Abcam; 07-662, Millipore; and 13826S, CST), Rplp1 (PA5-103540, Invitrogen), Lamp2 (PA1-655, Invitrogen), Ctsd (sc377299, Santa Cruz), Gapdh (2118, CST), cleaved-Casp3 (9661, CST), Map1lc3b-ii/Lc3b (a2775, CST), and Sqstm1/p62 (5114, CST) were used. Horseradish peroxidase–conjugated secondary antibodies recognizing mouse (sc-2954) and rabbit (sc-2955) immunoglobulin Gs (IgGs) were purchased from Santa Cruz Biotechnology. Blots were observed on Image Lab software (BioRad) and densitometric analysis was performed using ImageJ software (NIH, Bethesda, MD, USA) normalized to GAPDH as loading controls.

*Immunoprecipitation of puromycin-labeled proteins*

Protein immunoprecipitation was performed using Dynabeads™ Protein G for Immunoprecipitation (Invitrogen) as per manufacturer’s protocol. Non-denaturing lysis buffer (Abcam) was used to prepare tissue homogenate. Puromycin (1:5,000 dilution; MABE343, Merck), or 4 µg normal rabbit IgG as control (12-370, Sigma-Aldrich) was used for pull-down assay. Western blot analysis for the detection of pulled down proteins was performed as described above.

*Chromatin Immuno-Precipitation (ChIP)-qPCR in AML12 cells and ChIPseq in liver tissues*

ChIP-qPCR was performed as described previously [13, 15]. For Esrra binding on the *Esrra* gene, primer pairs [ATGCATGGTCCCA- GAGTCAG (forward) and CTGGTTTGCGAGTTCCTCAA (reverse)] were used to amplify −629 to −498 region on *Esrra* promoters in AML12 cells. For Polr2a binding, a primer pair [ATTAGCATAGGGCACCTGGC (forward) and CGACCAC- CGTGGCTGAC (reverse)] was used to amplify −69 to +11 region in AML12 cells. However, on the *Rplp1* gene promoter, for Esrra binding, primer pairs [AGCTTCTTTGTGGTCCTGAGATT (forward) and ACCCCTCAGGTTAGGGTACA (reverse) were used to amplify −988 to −837 region on the *Rplp1* promoter in AML12 cells. For Polr2a binding on *Rplp1* gene promoter, a primer pair [CAATCGCACCGGAAGTCGAA (forward) and GACCAGTCCACCTATATACGCC (reverse)] were used to amplify −140 to +18 region in AML12 cells.

ChIPseq methodology is provided elsewhere [12].

*Polysome Profiling in vitro Using Sucrose Gradients*

AML12 cells were cultured with BSA (0.5% as control), PA (0.5 mM for 24 h) or PA with si*Esrra*, washed with cold PBS and subsequently lysed on ice using a lysis buffer containing Tris-Cl (pH 7.4, 20mM), NaCl (150mM), MgCl2 (5mM), NP40 (1%), DTT (1µM), Cycloheximide (100µg/ml), and cOmplete™ EDTA-free Protease Inhibitor (1X). The lysate was then homogenized by passing through a 27G needle five times. Cellular debris was separated by centrifuging at 20,000xg for 10 minutes at 4°C. Following this, the lysate was loaded onto a 10-50% sucrose gradient, which had been previously prepared with the Gradient Master. Polysome profiling was subsequently conducted, and fractions were collected using the Biocomp fractionator, in tandem with TRIAX. The data was plotted using GraphPad PRISM software.

*Polysome Profiling in vivo (liver) Using Sucrose Gradients*

Sucrose gradient was prepared in polypropylene tube (344059, Beckman Coulter) by layering 50% of sucrose under 10% sucrose in 25 mM Tris, pH 7.5, 150 mM NaCl, 5 mM MgCl2 buffer solution. The sucrose gradient was formed using BioComp gradient master following manufacturer’s instruction. 100 mg of fresh liver was homogenized by passage through a micro mincer (294103, BIOSPEC) and the homogenate was lysed in 1ml of polysome lysis buffer containing 25 mM Tris, pH 7.5, 150 mM NaCl, 5 mM MgCl2 and 1% NP40 supplemented with 1X cOmplete™ Mini without EDTA, 0.75mg/ml heparin, and 1mM DTT. Tissue lysate was further homogenized by passage through a 25G syringe needle 5 times and incubated on ice for 10 minutes. Cell debris was removed by centrifuging the lysate at 20,000 xg for 10 minutes at 4oC. 250ul of the supernatant was overlaid on the 10-50% sucrose gradient which was subjected for ultra-centrifugation at 36,000 r.p.m for 2 hours at 4oC, with acceleration speed 7 and deceleration speed 1, using ultracentrifuge with rotor SW41Ti. The sucrose gradient was fractionated using BioComp piston gradient fractionator. Polysome profile was analyzed with TRiAX software (BioComp).

*Label-free quantitative proteomic analysis*

Label-free quantitative analysis of proteins, recovered from pooled triplicates each group, by LC-MS/MS was performed by NonovogeneAIT (Singapore) as described below.

Protein Quality Test: BSA standard protein solution was prepared according to the instructions of Bradford protein quantitative kit, with gradient concentration ranged from 0 to 0.5 g/L. BSA standard protein solutions and sample solutions with different dilution multiples were added into 96-well plate to fill up the volume to 20 µL, respectively. Each gradient was repeated three times. The plate was added 180 μL G250 dye solution quickly and placed at room temperature for 5 minutes, the absorbance at 595 nm was detected. The standard curve was drawn with the absorbance of standard protein solution and the protein concentration of the sample was calculated. 20 μg of the protein sample was loaded to 12% SDS-PAGE gel electrophoresis, wherein the concentrated gel was performed at 80 V for 20 min, and the separation gel was performed at 120 V for 90 min. The gel was stained by coomassie brilliant blue R-250 and decolored until the bands were visualized clearly.

Trypsin treatment: 120 μg of each protein sample was taken and the volume was made up to 100 μL with dissolution buffer, 1.5 μg trypsin and 500 μL of 100 mM TEAB buffer were added, sample was mixed and digested at 37 °C for 4 h. Andt hen,1.5 μg trypsin and CaCl2 were added, sample was digested overnight. Formic acid was mixed with digested sample, adjusted pH under 3, and centrifuged at 12000 g for 5 min at room temperature. The supernatant was slowly loaded to the C18 desalting column, washed with washing buffer (0.1% formic acid, 3% acetonitrile) 3 times, then eluted by some elution buffer (0.1% formic acid, 70% acetonitrile). The eluents of each sample were combined and lyophilized.

LC-MS/MS Analysis: Mobile phase A (100% water, 0.1% formic acid) and B solution (80% acetonitrile, 0.1% formic acid) were prepared. The lyophilized powder was dissolved in 10 μL of solution A, centrifuged at 14,000 g for 20 min at 4 °C, and 1 μg of the supernatant was injected into a home-made C18 Nano-Trap column (2 cm×75 μm, 3 μm). Peptides were separated in a home-made analytical column (15 cm×150 μm, 1.9 μm), using a linear gradient elution. The separated peptides were analyzed by Q Exactive HF-X mass spectrometer (Thermo Fisher), with ion source of Nanospray Flex™（ESI, spray voltage of 2.3 kV and ion transport capillary temperature of 320°C. Full scan range from m/z 350 to 1500 with resolution of 60000 (at m/z 200), an automatic gain control (AGC) target value was 3×106 and a maximum ion injection time was 20 ms. The top 40 precursors of the highest abundant in the full scan were selected and fragmented by higher energy collisional dissociation (HCD) and analyzed in MS/MS, where resolution was 15000 (at m/z200), the automatic gain control (AGC) target value was 1×105, the maximum ion injection time was 45 ms, a normalized collision energy was set as 27%, an intensity threshold was 2.2×104, and the dynamic exclusion parameter was 20 s. The raw data of MS detection was named as “.raw”.

The identification and quantitation of protein: All resulting spectra were searched against Mus_musculus_uniprot_2019.01.18.fasta (85165 sequences) database by the search engines: Proteome Discoverer 2.2 (PD 2.2, Thermo). The search parameters are set as follows: mass tolerance for precursor ion was 10 ppm and mass tolerance for product ion was 0.02 Da. Carbamidomethyl was specified as fixed modifications, Oxidation of methionine (M) was specified as dynamic modification, and acetylation was specified as N-Terminal modification in PD 2.2. A maximum of 2 missed cleavage sites were allowed. In order to improve the quality of analysis results, the software PD 2.2 further filtered the retrieval results: Peptide Spectrum Matches (PSMs) with a credibility of more than 99% was identified PSMs. The identified protein contains at least 1 unique peptide. The identified PSMs and protein were retained and performed with FDR no more than 1.0%. The protein quantitation results were statistically analyzed by T-test. The proteins whose quantitation significantly different between experimental and control groups, (p < 0.05 and |log2FC| >= 2 (ratio >= 4 or ratio <= 0.25 [fold change, FC]), were defined as differentially expressed proteins (DEP).

The functional analysis of protein and DEP: DisGeNET, Reactome 2022 and KEGG (Kyoto Encyclopedia of Genes and Genomes) 2021 were used to analyze the protein family and pathway on EnrichR platform (the Ma'ayan Lab, NY, USA) [17-19].

*Statistical analysis*

Individual culture experiments were performed in triplicate and repeated at least three times independently using matched controls; the data were pooled (n=3-5/group), and statistical analysis was performed. Animal studies were performed as per the approved IACUC protocol, and the statistical analysis were performed (n=4-5/group) as per the biostatistician’s advice. Results are expressed as mean ± SD for all *in vitro* and *in vivo* experiments. Normality of the data was analyzed by Shapiro-Wilk test, then a parametric analysis was performed using unpaired student’s t-test, one-way ANOVA or two-way ANOVA followed by Tukey’s multiple-comparisons test, wherever applicable. For any non-normal dataset, non-parametric Kruskal-Wallis test for ANOVA followed by Dunn's multiple comparisons test was performed. The statistical significance of differences was assessed as *P<0.05; **P<0.01; ***P<0.001; ****P<0.0001). All statistical tests were performed using Prism 10 for Mac OS X (GraphPad Software).

**Supplementary Tables (1):** *Supplementary tables for proteomic disease discovery and molecular pathways analysis can be found in one excel file submitted separately as Supplementary_information_Proteome_Dataset_analysis.xlsx.*

**Supplementary Table (2):** Patient details

| **Stage** | **Histology**  **(fibrosis/ cirrhosis)** | **BMI** | **Weight (Kg)** | **Age** | **Sex** | **Drinker/ non-drinker** | **Ethnicity** |
| --- | --- | --- | --- | --- | --- | --- | --- |
| Normal | N | NA | NA | 56 | M | N | Chinese |
|  | N | NA | NA | 39 | F | NA | Others |
|  | N | NA | NA | 76 | M | N | Malay |
|  | N | 24.83 | 61.2 | 73 | F | N | Chinese |
|  | N | NA | NA | 52 | M | N | Chinese |
|  | N | 24.6 | 71 | 61 | M | N | Chinese |
|  | N | 20.8 | 55.2 | 50 | M | N | Others |
|  | N | 28.1 | 81.1 | 70 | M | N | Chinese |
|  | N | 23.4 | 62.9 | 77 | M | N | Chinese |
|  | N | 19.38 | 56 | 60 | M | N | Others |
|  | N | 19.7 | 57 | 69 | M | N | Others |
| NAFLD | N | 23.1 | 64.5 | 59 | M | N | Chinese |
|  | N | 24.8 | 61 | 45 | M | N | Indian |
|  | N | NA | NA | 69 | M | N | Others |
|  | N | NA | NA | 44 | M | N | Chinese |
|  | N | 23.1 | 71.4 | 71 | M | N | Others |
|  | N | 26.4 | 65 | 70 | F | N | Malay |
|  | N | 24.7 | 63.9 | 68 | M | N | Chinese |
| NASH | N | 23.2 | 67 | 56 | M | N | Others |
|  | Fibrosis | 23.2 | 67.7 | 72 | M | N | Others |
|  | Fibrosis | 32.7 | 90.1 | 64 | M | N | Chinese |
|  | N | NA | NA | 55 | F | N | Malay |
|  | Fibrosis | 19.2 | 51.6 | 67 | M | N | Chinese |
|  | Fibrosis | 20.3 | 60.1 | 71 | M | N | Chinese |
|  | Cirrhotic | 23.1 | 71.4 | 71 | M | N | Others |
|  | Cirrhotic | 19.4 | 49.1 | 54 | M | N | Chinese |
|  | Cirrhotic | 29.69 | 84.8 | 53 | M | N | Others |

**N**, No; **NA**, data not available
